# Isolation and complete genome sequencing of Pectobacterium phage Jarilo, representing a novel genus of bacteriophages within the subfamily *Autographivirinae*

**DOI:** 10.1101/2020.01.16.909176

**Authors:** Julie Stenberg Pedersen, Alexander Byth Carstens, Amaru Miranda Djurhuus, Witold Kot, Lars Hestbjerg Hansen

## Abstract

*Pectobacterium carotovorum* is the causative agent of bacterial soft rot on various plant species. The use of phages for plant disease control have gained increased awareness over the past years. We here describe the isolation and characterization of Pectobacterium phage Jarilo, representing a novel genus of bacteriophages within the subfamily *Autographivirinae*. Jarilo possesses a double-stranded DNA genome of 40557 bp with a G+C% content of 50.08% and 50 predicted open reading frames (ORFs). Gene synteny and products seem to be somewhat conserved between Pectobacterium phage Jarilo and Enterobacteria phage T7, but limited nucleotide similarity is found between Jarilo and other phages within the subfamily *Autographivirinae.* We propose Pectobacterium phage Jarilo as the first member of a new genus of bacteriophages within the subfamily *Autographivirinae.*

## Introduction

*Pectobacterium carotovorum* is part of the family *Pectobacteriaceae*, which members include important bacterial plant pathogens involved in a wide range of diseases[1]. *Pectobacteriaceae* are Gram-negative, facultative anaerobes, that are non-spore forming and produce extracellular enzymes, involved in their pathogenicity[2]. *P. carotovorum* (formerly *Erwinia corotovora*) is able to form soft rot disease symptoms in various plant species. In potatoes *P. carotovorum* is the causative agent of black leg and tuber rot during storage, affecting postharvest loss[3,4]. Control mechanism of bacterial soft rot is limited and is primarily based on sanitary conditions and good agricultural practice[5]. Due to limited control mechanism, increased attention have been directed towards biological control agents[6]. Phages have been proposed as promising biocontrol agents towards *Pectobacteriaceae* in several studies[7–12]. In this report, we describe the isolation and characterization of Pectobacterium phage Jarilo, representing a new genus of bacteriophages within the subfamily *Autographivirinae*.

## Materials and methods

Jarilo was isolated from an organic waste sample as described previously[12], using *Pectobacterium carotovorum* DSM 30170 as host. Prior to isolation virus particles in the organic waste sample was concentrated by polyethylene glycol precipitation, as described elsewhere[13]. Phage DNA was isolated and sequencing libraries prepared using direct plaque sequencing protocol[14], with the modifications described previously[15]. Reads from sequencing were trimmed and *de novo* assembled using CLC Genomic Workbench (10.1.1). Open reading frame prediction were automatically executed using DNA Master (version 5.0.2), with GeneMark[16] and Glimmer[17] as gene caller, and were corrected manually. Putative gene functions were manually curated using BLASTp[18] and HHpred[19] with the databases Pfam (version 32.0), SCOP70 (version 1.7.5) and pdb70. tRNA sequences were searched for using tRNAscan-SE[20].

## Results

Pectobacterium phage Jarilo represents a new genus within the subfamily *Autographivirinae*, with limited DNA sequence similarity (<50%) with other phages within the same subfamily (Fig. 1)[21]. Pectobacterium phage Jarilo has a double-stranded DNA genome of 40557 bp with a G+C% content of 50.08%. In the genome there is 50 predicted open reading frames (ORFs), whereas the majority (30/50) could be assigned with a putative function due to homology with other known phages within the subfamily *Autographivirinae*. No tRNAs were detected.

**Figure 1.**
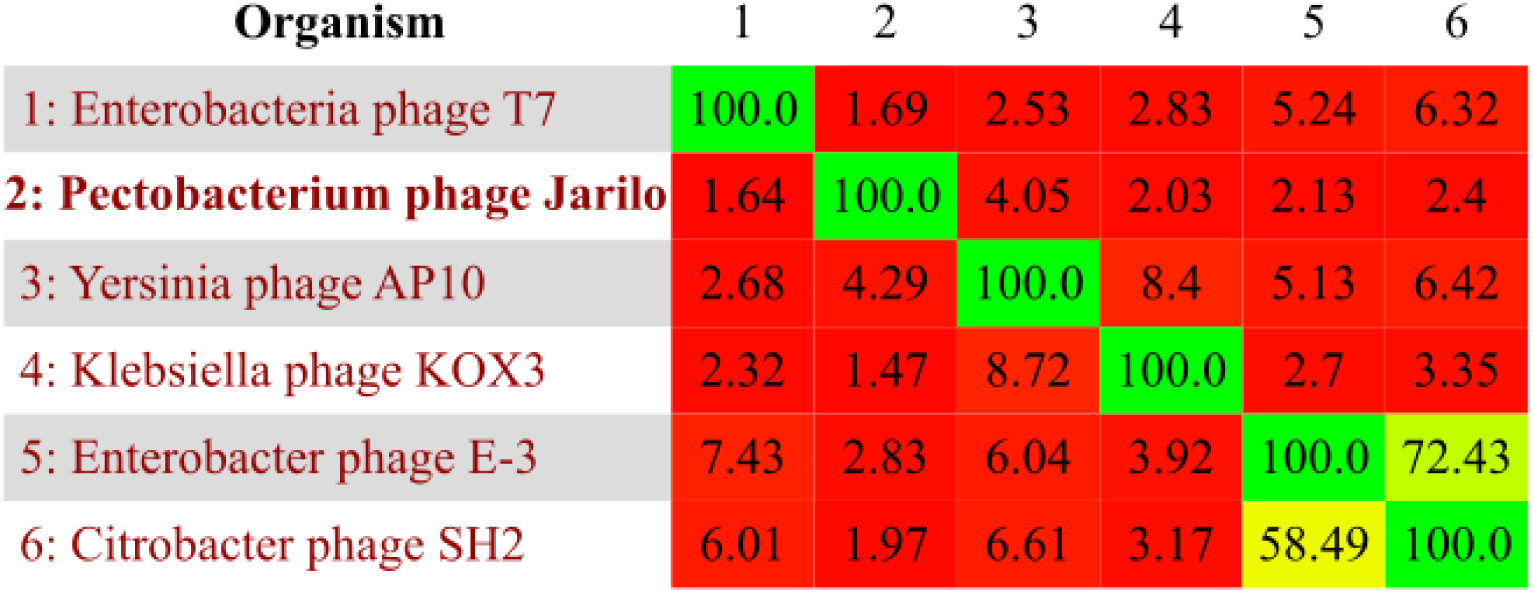
Pectobacterium phage Jarilo possesses limited DNA sequence similarity with other known *Autographivirinae* phages. All vs. all nucleotide comparison between six genomes of bacteriophages within the subfamily *Autographivirinae*, using the Gegenees software (fragment size: 500 bp, step size: 500 bp)[27]. Phages showing highest scores, using blastn[18], against the genome of Jarilo were chosen together with phage T7 as a reference genome. Colors indicate similarity between phages, from similar (green) to less similar (red). Similarity scores are based on blastn. Following phage genomes were used (with genbank accession numbers); Yersinia phage AP10 (KT852574.1), Klebsiella phage KOX3 (MN101216.1), Enterobacter phage E-3 (KP791806.1), Citrobacter phage SH2 (KU687348.1), Enterobacteria phage T7 (V01146.1).

Based on bioinformatics predictions, ORFs were categorized into five putative gene functions; hypothetical protein (grey), DNA replication and metabolism (red), lysis proteins (yellow), others/unknown (green) and morphogenesis (blue) (Fig. 2). The gene synteny of the genome of Jarilo is conserved among phages within the subfamily *Autographivirinae*, as well as many of the structural genes (Fig. 2). An insertion of a putative endonuclease VII appear to have happened within the putative DNA polymerase, in the genome of Jarilo. This insertion separates the DNA polymerase into two ORFs, both ORFs share high sequence similarity with the DNA polymerase of related phages (Fig. 2). Yersinia phage YpP-G, also part of the *Autographivirinae* subfamily, encodes a endonuclease VII, which gives the highest score against the putative endonuclease VII encoded by Jarilo, using blastp (Suppl. fig. 1)[18,22]. YpP-G display a similar gene arrangement with an endonuclease VII within the gene encoding the DNA polymerase (Suppl. fig. 1). Furthermore is the same phenomenon described in another phage within the subfamily *Autographivirinae*, Pf-WMP3[23]. A endonuclease VII insertion into the DNA polymerase have been proven as part of a self-splicing intron[24], but have not been demonstrated as part of a selfsplicing intron within the *Autographivirinae* subfamily. The gene encoding the major capsid protein as well as the gene following, categorized as a hypothetical gene, in Jarilo, resembles the capsid gene in T7 with some similarity. The capsid gene in T7 has two products, a major (10A) and a minor (10B), produced by frameshifting into the −1 frame near the end of 10A[25]. The gene encoding head to tail joining protein appear to be divided in two (Fig. 2), which could also be a result of frameshifting, as in the capsid protein of T7, or due to a nonsense mutation. Genes categorized as morphogenesis, in Jarilo, are conserved among phages within the subfamily *Autographivirinae*, however is the tail-fiber protein conserved to a lesser extent, which could represent different receptor preferences (Fig. 2). Interestingly, Jarilo, T7 and AP10 shares a conserved region at the terminal of the genome, not representing an ORF (Fig. 2). Like other phages within the subfamily *Autographivirinae*, Jarilo encodes a RNA polymerase[26] and do not encode any detectable integrase.

**Figure 2.**
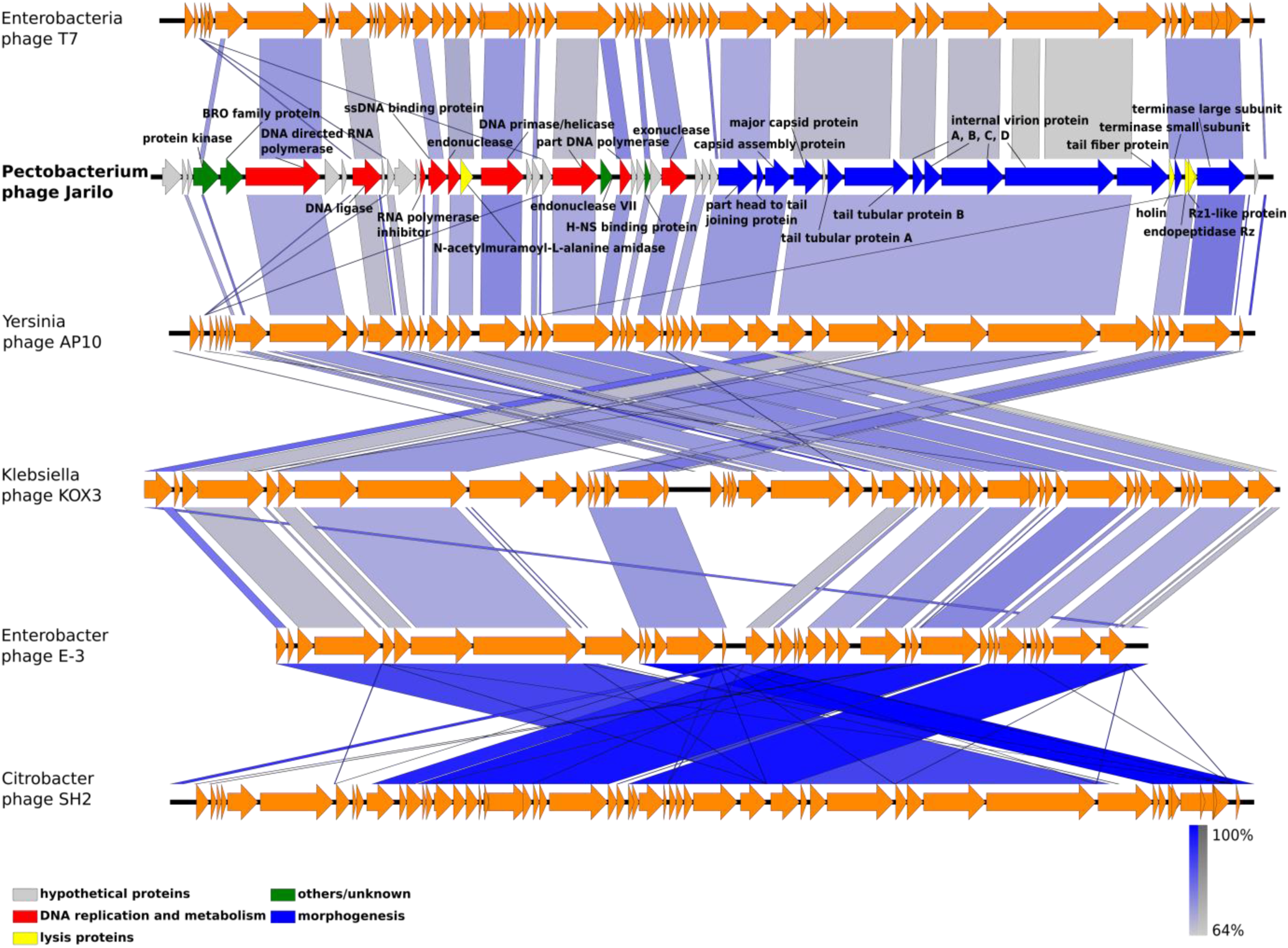
Comparative genomics reveals conserved gene synteny among phages within the subfamily *Autographivirinae*, as wells as a high degree of sequence homology among structural proteins. Comparative genomics of six genomes of bacteriophages within the subfamily *Autographivirinae* using blastn in the Easyfig software (version 2.2.3)[28]. Phages showing highest scores, using blastn[18], against the genome of Jarilo were chosen together with phage T7 as a reference genome. Putative gene functions of the genome of Jarilo were divided into five categories; hypothetical proteins (grey), DNA replication and metabolism (red), lysis proteins (yellow), others/unknown (green) and morphogenesis (blue). Following phage genomes were used (with Genbank accession numbers); Yersinia phage AP10 (KT852574.1), Klebsiella phage KOX3 (MN101216.1), Enterobacter phage E-3 (KP791806.1), Citrobacter phage SH2 (KU687348.1), Enterobacteria phage T7 (V01146.1).

Based on the findings we propose Pectobacterium phage Jarilo as the first representation of a new genus of bacteriophages belonging to the subfamily *Autographivirinae*, family *Podoviridae*, order *Caudovirales*.

## Nucleotide sequence accession number

The GenBank accession number for Pectobacterium phage Jarilo is MH059637.1.

