## Supplementary figure 1 for "Isolation and complete genome sequencing of Pectobacterium phage Jarilo, representing a novel genus of bacteriophages within the subfamily *Autographivirinae*"

### Supplementary figures:

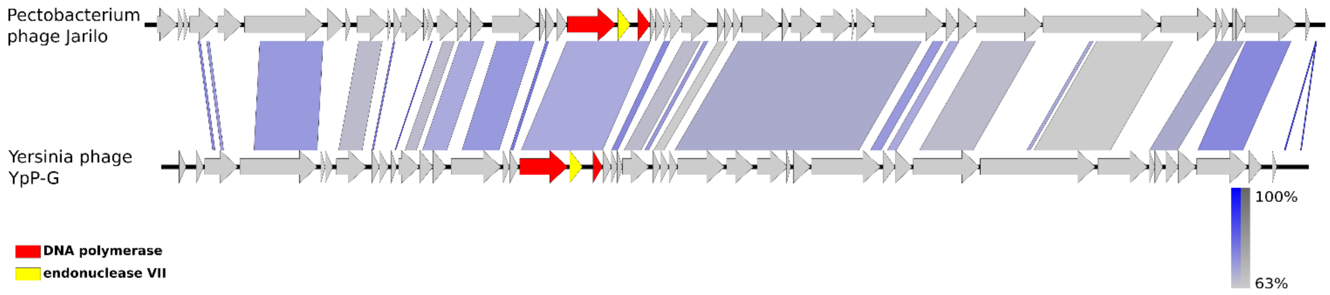


**Suppl. fig. 1. The DNA polymerase in two ORFs divided by a endonuclease VII, in Pectobacterium phage Jarilo and Yersinia phage YpP-G**

Comparative genomics of the genomes of Jarilo and YpP-G (GenBank: JQ965702.1)[22] using blastn in the Easyfig software (version 2.2.3)[28], to visualize conservation within the genes encoding DNA polymerase (red) as well as the endonuclease VII within this gene (yellow).
